# A homogenization approach for spatial cytokine distributions in immune-cell communication

**DOI:** 10.64898/2026.03.31.715485

**Authors:** Lisa Li, Lorena Pohl, Andreas Hutloff, Barbara Niethammer, Kevin Thurley

**Affiliations:** Biomathematics Division, Institute of Experimental Oncology, University Hospital Bonn, Germany; Institute for Applied Mathematics, University of Bonn, Germany; Institute of Medical Immunology, University Hospital Schleswig-Holstein Campus Kiel, Germany; Bonn Center for Mathematical Life Sciences, University of Bonn, Germany

## Abstract

Cytokine-mediated communication is a central mechanism by which immune cells coordinate activation, differentiation and proliferation. While mechanistic reaction-diffusion models provide detailed descriptions of cytokine secretion and uptake at the cellular scale, their computational cost limits their applicability to large and densely packed cell populations. Previously employed approximations of cytokine diffusion fields rely on assumptions that neglect the influence of cellular geometry and volume exclusion. In this work, we study a macroscopic description of cytokine diffusion and reaction dynamics based on homogenization techniques, rigorously linking microscopic reaction–diffusion formulations to effective continuum models. The resulting homogenized equations replace discrete responder cells with a continuous density, while retaining essential features of cellular uptake and excluded-volume effects. Further, we show that in regimes with approximate radial symmetry, classical Yukawa-type solutions emerge as limiting cases of the homogenized model, provided appropriate correction factors are included. Overall, our approach allows efficient multiscale modeling of cytokine signaling in complex immune-cell environments.

## 1 Introduction

CD4^+^ T helper (Th) cells orchestrate both innate and adaptive immune responses and are therefore essential for effective cellular immunity against pathogens. Cell-cell communication mediated by soluble cytokines plays a central role in directing differentiation of naive Th cells into distinct effector subsets, thus mounting an immune response tailored to a specific pathogenic threat (Lin and Leonard, 2019; O’Shea and Paul, 2010). On the other hand, Th cells are defined by a characteristic profile of secreted cytokines, for example T helper type 1 (Th1) cells are characterized by expression of interferon-*γ* (IFN-γ), enabling them to effectively combat intracellular bacteria. Another specialized subset, follicular T helper (Tfh) cells, provides help to B cells and is an important source of interleukin(IL)-21, and is therefore indispensable for the generation of high-affinity, pathogen-neutralizing antibodies (Ritzau-Jost and Hutloff, 2021). An inappropriate Th subset response to a given pathogen can lead not only to inefficient pathogen clearance but also to severe immunopathology and, ultimately, death of the host. Further, IFN-*γ* and IL-21 promote T-cell differentiation into the corresponding effector populations (Th1 or Tfh cells), by binding to their cognate receptors on the T-cell surface and activating intracellular signaling pathways (Ritzau-Jost and Hutloff, 2021). Hence, cytokine signals form the basis of complex cell-cell interaction networks that are essential for the regulation of immune-responses, and a quantitative understanding of those networks is only beginning to emerge (Morel et al., 2017; Steinheuer et al., 2025).

Paracrine cytokine signaling amongst tissue cells naturally imposes the question of spatial patterning. The existence of spatial cytokine inhomogeneities within a physiological parameter regime was predicted and systematically analyzed by model simulations (Brunner et al., 2024; Busse et al., 2010; Fuhrmann et al., 2016; Thurley et al., 2015), and indeed, subsequent experimental work supported tunable inhomogeneities in cytokine signals within lymphoid tissue (Oyler-Yaniv et al., 2017; Wong et al., 2021).

Specifically, a reaction-diffusion (RD) model to describe the spatial distribution of a cytokine species *c* has been set up as follows (Brunner et al., 2024):

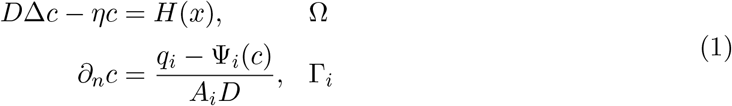

where Ω denotes the extracellular space, Γ_*i*_ the cell surfaces with area *A*_*i*_, and *η* and *D* cytokine degradation rate and the diffusion constant, respectively. The source term *H*(*x*) represents system-wide cytokine secretion and uptake in the extracellular space, for instance by the extracellular matrix or other cell types. The cell-surface mediated cytokine secretion rate is denoted by *q*_*i*_, and Ψ(*c*) represents cellular uptake of cytokine molecules. Of note, Equation (1) is based on a quasi-steady-state assumption for the cytokine concentration, since the time-scale of diffusion across a typical cell-cell distance (seconds) is much faster than the time-scale of biochemical reactions (minutes to hours). Further, Ψ(*c*) is taken as a non-linear, saturating function, since responder cells can only express a limited number of cytokine receptor molecules and thus have a limited uptake capacity. Equation (1) provides a mechanistic description of cytokine secretion and uptake. However, it can be computationally demanding, especially when considering large cell-populations with complex geometries and communicating by several cytokine species. On the other hand, available explicit descriptions of cytokine fields (Olimpio et al., 2018; Oyler-Yaniv et al., 2017) rely on the Yukawa-potential, which assumes a homogeneous receptor density and implicitly neglects cell volumes.

Here, we present a rigorous derivation of a macroscopic model for the cytokine concentration along with a numerical study of its validity and a computational framework for modeling cell-cell interactions considering spatial cytokine gradients. We use the method of homogenization, a mathematical technique used to derive effective macroscopic properties of materials or systems that exhibit fine-scale heterogeneity, for example materials containing many small holes (Cioranescu and Donato, 1988; Cioranescu and Murat, 2018; Cioranescu and Paulin, 1979; Conca and Donato, 1988). These techniques have also been used to derive effective equations for ligands in time-dependent systems with uptake depending on the concentration of surface receptors (Marciniak-Czochra and Ptashnyk, 2008; Ptashnyk and Venkataraman, 2020). Here we consider a simplified model where the limited number of receptor molecules is expressed through a nonlinear saturated uptake function.

The paper is structured as follows: After deriving the macroscopic equations (Section 2), we focus on special cases and retrieve a modified Yukawa potential with correction factors accounting for excluded-volume effects (Section 3). Furthermore, we investigate the conditions and parameter regimes under which the homogenized model provides a good approximation of the precise model, and we apply the derived methodology to cytokine-mediated decision-making during Th cell differentiation (Section 4).

## 2 Homogenization approach to spatial cytokine gradients

To derive a macroscopic formulation of spatio-temporal cytokine dynamics in the extracellular domain, consider a single cytokine-secreting cell surrounded by non-secreting responder cells. To specify Equation (1), we imposed saturating uptake functions of the form

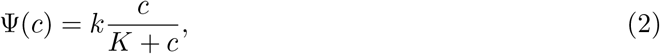

where *k* is the maximal uptake rate and *K* is the half-saturation constant.

The next step is to non-dimensionalize Equations (1)-(2) for the cytokine diffusion field. Note that the cytokine concentration *c* is given in units of molecules per volume. We introduced dimensionless variables by means of *x* := *x/d, u*(*x*) := *c/K, h*(*x*) := *H*(*x*)*d*^2^*/*(*DK*), *Ψ*(*u*) := *u/*(*u* + 1), yielding

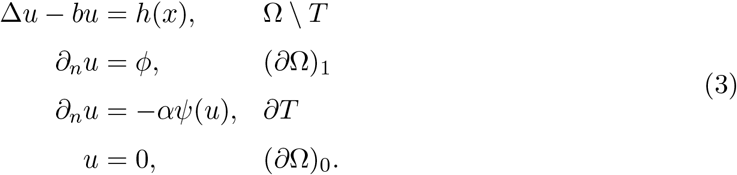

Here, *d* denotes the cell-cell distance, and we identified the dimensionless parameters *b* = *ηd*^2^*/D, ϕ* = *qd/*(*ADK*), *α* = *dk/*(*ADK*). Throughout, we assumed *u*(*x*), *x* ∈ Ω to be in quasi-steady-state, and further that cytokine secretion and uptake are homogeneous across each cell surface. Setting Ω as the spatial domain, where the secreting cell is excluded, and *T* as the collection of all responder cells, the diffusion area takes the form Ω \ *T*. The boundary ∂Ω is split into the external boundary (∂Ω)_0_ and the secreting-cell boundary (∂Ω)_1_ (Figure 1a).

**Figure 1.**
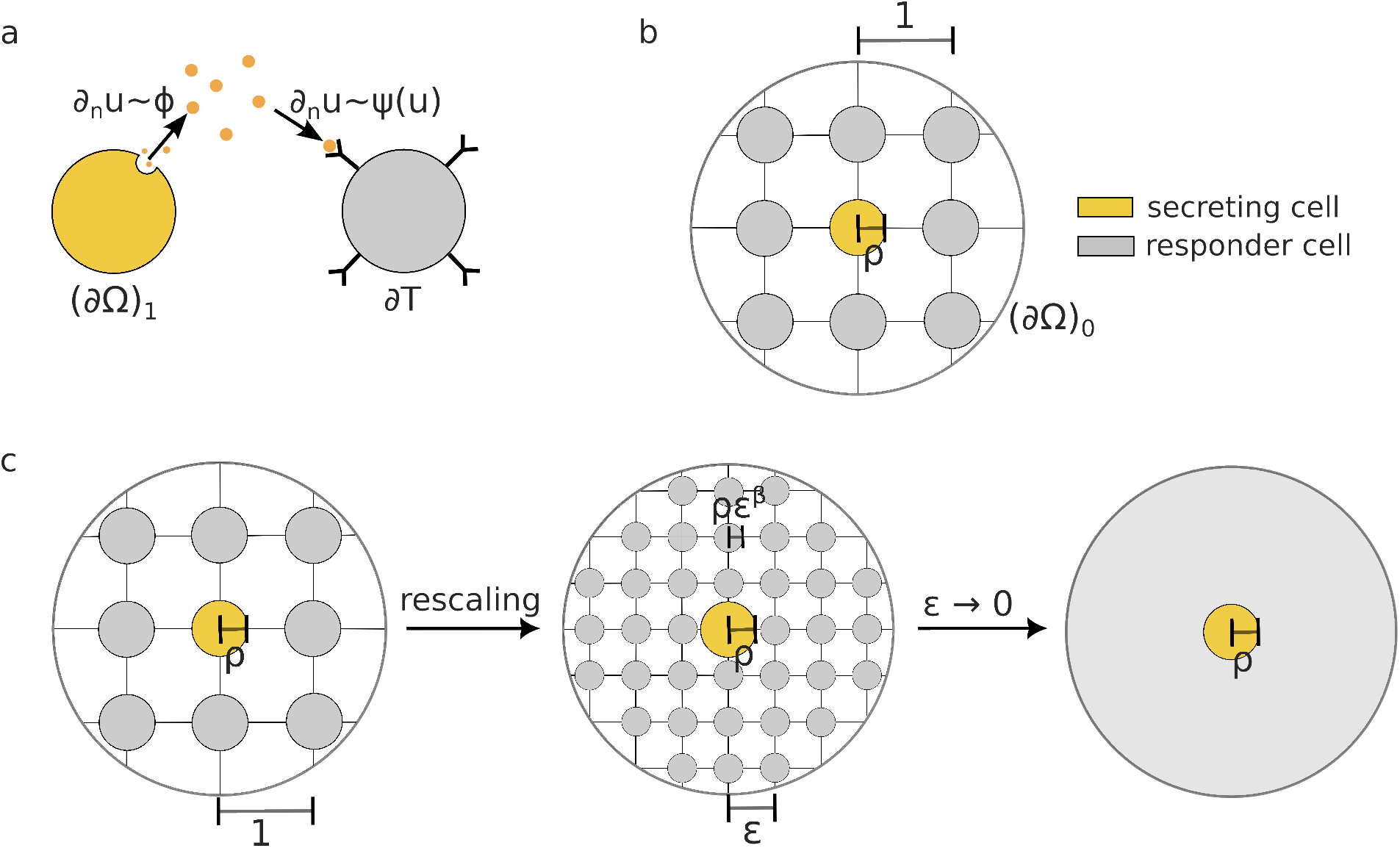
Schematic representation of the cytokine signaling model and the homogenization method. (a) Model scheme. On the boundary of secreting cells (∂Ω)_1_, cytokine secretion is proportional to the secretion rate *ϕ* and on the boundary of responder cells ∂*T*, cytokine uptake is proportional to the uptake function *ψ*. (b) Geometric set-up. Responder cells are arranged on a cubic grid of size 1, surrounding one secreting cell at the center. (c) Homogenization method. By introducing a scaling factor *ϵ*, both the responder cell radius and the cell-cell distance are scaled to 0 in the limit *ϵ* → 0.

Intuitively, Equation (3) resembles a generalized screened Poisson equation

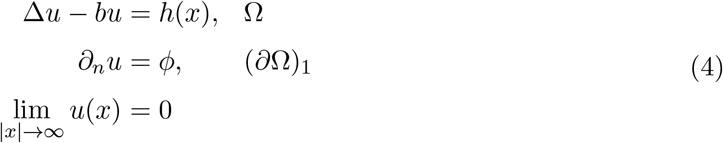

Hence, assuming radial symmetry and an uptake field *h*(*x*) = *α*_*total*_*u* with total uptake rate *α*_total_, we expect the solution to be of Yukawa-type

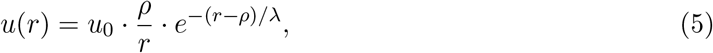

where the screening length 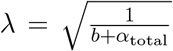can be identified with the spatial decay of the cytokine profile and *u*_0_ is a normalization constant (Oyler-Yaniv et al., 2017). However, the Yukawa derivation relies on strong assumptions such as radial symmetry and is furthermore incompatible with the non-linear uptake kinetics given in Equation (3). A more accurate analysis is therefore required to determine a generalized mean-field description. To address this, we applied a homogenization approach by rescaling the system and considering the limit of increasingly dense cell populations.

For this purpose, we defined the domain Ω = *B*_*R*_(0)\*B*_*ρ*_(0), with *R > ρ*. Within this domain, the responder cells are arranged on a three-dimensional lattice of grid size 1, each represented by a sphere of radius *ρ* centered at a lattice point (Figure 1b). Next, Equation (3) is rescaled as follows. The cell-cell distance, or more precisely, the size of the cubic grid covering Ω, on which the responder cells are placed, is set to *ϵ >* 0. Further, we introduced the scaling laws (Figure 1c)

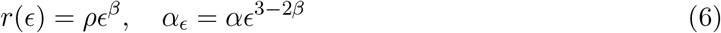

for some *β* ≥ 1. That is, the cell radius *r*(*ϵ*) of responder cells scales with the cell-cell distance by a power law, and the rescaled uptake function *α*_*ϵ*_ ensures a constant total uptake rate when *ϵ* → 0: while the number of responder cells scales with *ϵ*^−3^, the surface area of each responder cell decreases with *ϵ*^2*β*^. To define the domain of the rescaled cytokine diffusion problem, we dropped the responder cells that are not fully contained in Ω and denote the union of the remainder by *T*_*ϵ*_. This yields the perforated domain Ω_*ϵ*_ := Ω \ *T*_*ϵ*_. Altogether, we arrived at the rescaled PDE

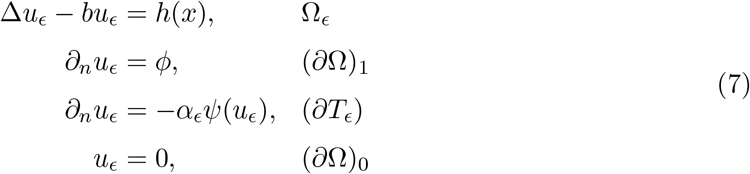

For given *ϵ >* 0, Equation (7) has a unique solution *u*_*ϵ*_ in *W*_*ϵ*_ := {*w* ∈ *W* ^1,2^(Ω_*ϵ*_) : *w* = 0 on (∂Ω)_0_}, which can be seen using the classical Lax-Milgram’s theorem.

To derive the homogenized cytokine field, we studied the limit of *u*_*ϵ*_ for *ϵ* → 0 in the Sobolev space *W* := {*w ∈ W* ^1,2^(Ω) : *w* = 0 on (∂Ω)_0_}. We found that the precise limit formulation of Equation (7) depends on the exponent *β* in Equation (6). We distinguish two cases: *β >* 1 and *β* = 1. For *β >* 1, in the homogenization process, the cell radius decreases faster than the cell-to-cell distance, and therefore the volume fraction of the responder cells becomes negligible. In this case, we have strong convergence of the characteristic function *χ*Ω_∈_ → 1 in *L*^2^(Ω). In contrast, for *β* = 1, the cell radius and the cell-to-cell distance have the same scaling behavior, and thus the volume fraction of responder cells remains constant. Assuming spherical cells for convenience, this leads to a constant excluded-volume fraction given by 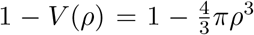 in the limit equation. More importantly, in this case, we only obtain weak convergence of the characteristic function 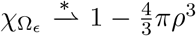. Overall, our analysis yields a modified diffusivity in the limit equation, and we derived the following

### Proposition 1

*Macroscopic formulation of cytokine diffusion*

A. *Let β* ∈ (1, 3) *and let u*_*ϵ*_ *be the unique solution of Equation* (7). *Then u*_*ϵ*_ ⇀ *u in W, where u is the unique weak solution of*

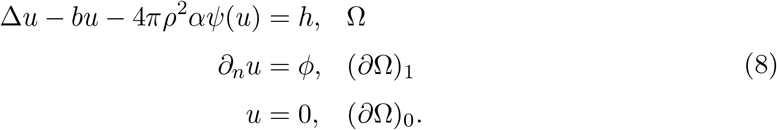
B. *Let β* = 1 *and let u*_*ϵ*_ *be the unique solution of Equation* (7). *Then u*_*ϵ*_ ⇀ *u in W, where u is the unique weak solution of*

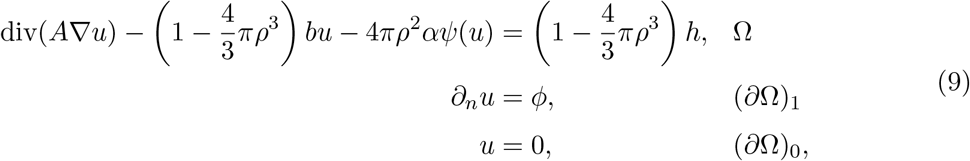

*where the Laplace modification matrix A* ∈ ℝ^3*×*3^ *is given by the following cell problem: For i* = 1, 2, 3, *let w*_*i*_ *be the solution of*

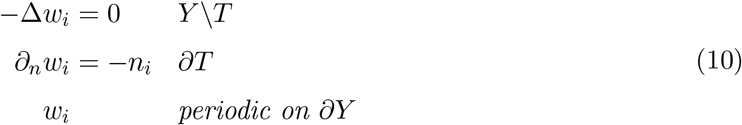

*on the reference cube* 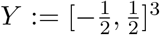 *with hole T* := *B*_*ρ*_(0). *Here, n*_*i*_ *denotes the i-th component of the normal vector at x* ∈ ∂*T*. *Then the i-th column of A is defined by*

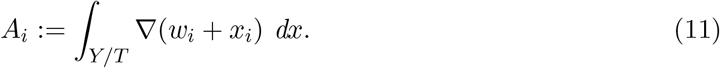

For the proof see Appendix, Section A.

### Remark 1

*In the case β* ≥ 3, *the size of the responder cells tends to 0 much faster than the cell-cell distance. Consequently, the rescaling required for the uptake function α*_*ϵ*_ *in order to obtain a nonzero contribution in the limit equation becomes very large. In fact, the rescaled uptake rate diverges so fast that it can no longer be controlled by the estimates used in the proof. That this case is not covered by Proposition 1 reflects the fact that, within our homogenization framework, responder cells whose size decreases too rapidly do not contribute to the homogenized limit equation*.

## 3 Special cases of the homogenized model

After deriving the limiting equations governing the homogenized model, we explored special cases where more explicit descriptions of the model can be derived. Such simplified formulations are particularly beneficial for applications, enabling more efficient numerical simulations. We started by investigating the cell problem (10) and the Laplace modification matrix *A* (11) under the assumption of ellipsoidal and spherical cells. Based on this analysis, we derived an ODE formulation of the limit equations under the assumption of radial symmetry, leading to an explicit solution in the case of linear uptake. Finally, assuming spherical cells, we numerically verified that the rescaled RD-system converges to the homogenized equation.

### 3.1 Laplace modification matrix *A* for ellipsoidal cells

In the case of *β* ∈ (1, 3), Proposition 1 directly yields the desired macroscopic formulation of the cytokine diffusion field in terms of Equation (8). However, the excluded-volume regime *β* = 1 leads to Equation (9), whose coefficients depend on the solution of the auxiliary boundary-value problem, Equation (10). For example, consider an ellipsoidal secreting cell *T* with semi-axes *ρ*_1_, *ρ*_2_, *ρ*_3_. In this case, the outward normal vector at a point *x* ∈ ∂*T* is given by 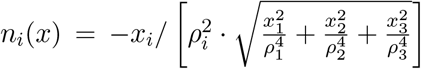. Since the solution *w*_1_ of Equation (10) is invariant under the transformation *φ*_*i*_ : ℝ^3^ → ℝ^3^, *x*_*i*_ ‘→ −*x*_*i*_ for *i* = 2, 3, we conclude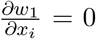. Applying the same argument to *w*_2_ and *w*_3_ implies that *A* = diag(*a*_1_, *a*_2_, *a*_3_) must be a diagonal matrix with *a*_1_, *a*_2_, *a*_3_ ∈ ℝ. In the case of a spherical cell *T*, the normal vector at *x* ∈ ∂*T* is 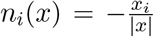The symmetric geometry implies that *w*_2_ and *w*_3_ can be obtained from *w*_1_ by relabelling the axes. Therefore, the diagonal entries of *A* must coincide, i.e. *a*_*i*_ = *a* ∈ ℝ for *i* = 1, 2, 3. Denoting the extracellular volume fraction by 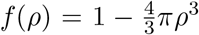, Equation (9) then simplifies to

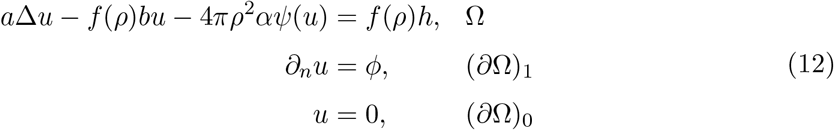

To illustrate those symmetry arguments, we numerically solved the cell problem, Equation (10), for a spherical cell with radius *ρ* and an ellipsoidal cell with axes *ρ*_1_ and *ρ*_2_ = *ρ*_3_ (Figure 2a), using a finite-element solver (Appendix B). From these solutions, we computed the matrix *A* using Equation (11). We observed that the off-diagonal entries of *A* converge to 0 as the mesh is refined. Furthermore, the diagonal entries converge to constants *a*_1_, *a*_2_, *a*_3_ ∈ R with *a*_2_ = *a*_3_ in the ellipsoidal case, reflecting the symmetry *ρ*_2_ = *ρ*_3_, and to the same value *a*_*sphere*_ in the spherical case (Figure 2b). Next, we evaluated the diagonal entries of *A* for varying cell geometries. For an ellipsoidal cell with varying axis ratio *ρ*_1_*/ρ*_2_ and fixed volume, we observed that *a*_1_ and *a*_2_ = *a*_3_ converge to the same value for *ρ*_1_*/ρ*_2_ → 1 (Figure 2c-d, upper plots). The diagonal entries further depend on the cell volume, in both the spherical and ellipsoidal case. For vanishing cell volume, i.e. *ρ* → 0, the diagonal entries and the extracellular volume factor *f*(*ρ*) converge to 1 (Figure 2c middle and lower panel, 2d lower plot).

**Figure 2.**
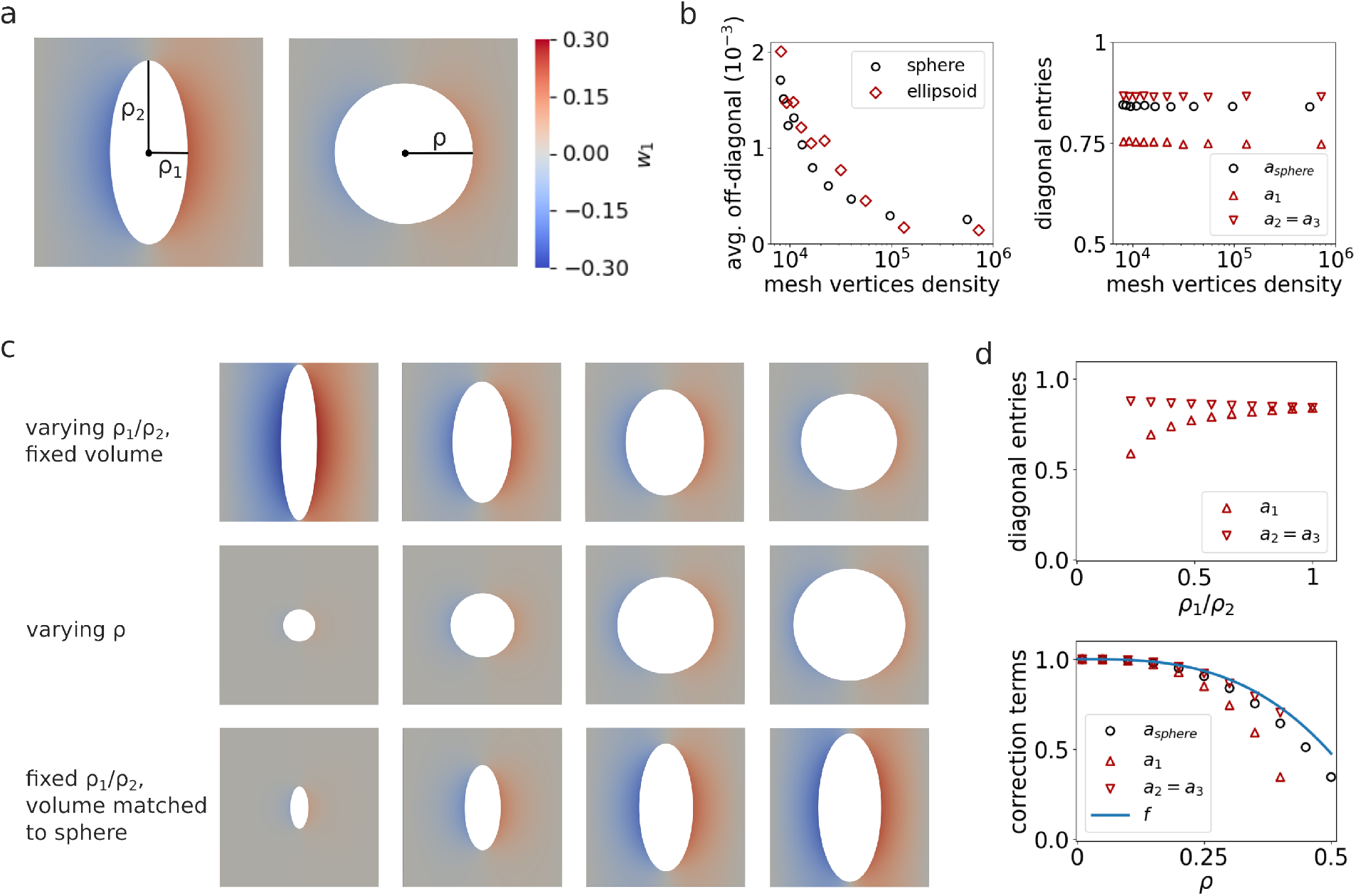
Simulation of the cell problem (10) and evaluation of the Laplace modification matrix *A* for various cell geometries. (a) Simulation of *w*_1_ in the case of an ellipsoidal cell with axes *ρ*_1_, *ρ*_2_, *ρ*_3_ where *ρ*_2_ = *ρ*_3_ and a spherical cell with radius *ρ*. (b) Evaluation of *A* under increasing mesh vertices density (number of mesh vertices divided by the volume of *Y* \ *T*). Left plot shows the average off-diagonal entries in the ellipsoidal and spherical case. Right plot shows the diagonal entries *a*_1_, *a*_2_, *a*_3_, with *a*_2_ = *a*_3_ in the ellipsoidal case, and the diagonal entry *a*_*sphere*_ in the spherical case. (c) Simulation of *w*_1_ for different cell geometries: an ellipsoidal cell with varying axis ratio *ρ*_1_*/ρ*_2_ assuming *ρ*_2_ = *ρ*_3_ and fixed volume (upper panel), a spherical cell with varying radius *ρ* (middle panel), and an ellipsoidal cell with fixed ratio *ρ*_1_*/ρ*_2_ and varying volume (lower panel). (d) Evaluation of the diagonal entries of *A* for the different scans visualized in (c). Lower plot shows the degradation correction term *f*(*ρ*) in addition.

Altogether, we found that the homogenized limit equation for *β* = 1 has the same structural form as for *β* ∈ (1, 3), but with excluded-volume correction terms. Specifically, the finite volume fraction of responder cells introduces a factor *f*(*ρ*) in the degradation and source term, and modifies diffusion through the coefficient *a*(*ρ*). In this sense, the excluded-volume limit does not recover the classical screened Poisson Equation (4), but rather a modified version with effective diffusion and degradation.

### 3.2 ODE formulation in the case of spherical cells

In a radial-symmetric setting, the homogenized descriptions of cytokine distribution lead to simplified formulations that enable more efficient simulations. To derive these, consider the case of spherical cells and *h* = 0. Then the solutions *u*(*x*) of the homogenized cytokine PDE (8) and (9) only depend on the radial coordinate, i.e. *u*(*x*) = *u*(*r*). Furthermore, since the Laplace operator in spherical coordinates simplifies to 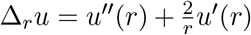, Equation (12) simplifies to the following ODE formulation:

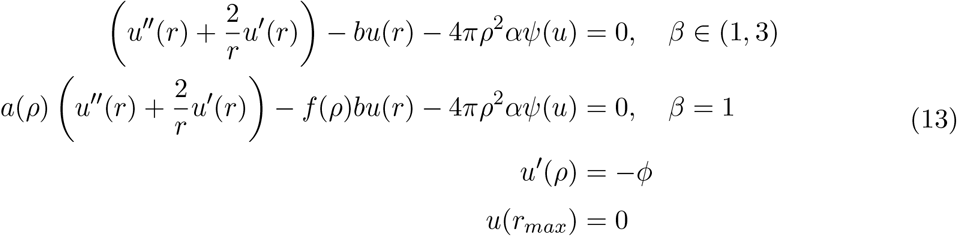

with the excluded-volume correction terms *f*(*ρ*) and *a*(*ρ*) defined above (Figure 2d, Equation 12).

In the case of linear uptake, i.e. *Ψ*(*u*) = *u*, and unbounded domain, i.e. lim_*r*→∞_ *u*(*r*) = 0, the ODE (13) yields the explicit solution

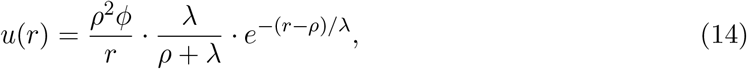

with signaling range

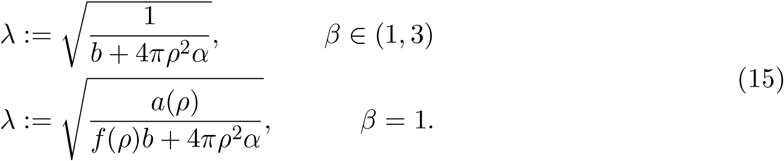

In particular, in the case *β* ∈ (1, 3) where the cell-volume is neglected in the limit of *ϵ* → 0, the Yukawa potential (5) is recovered for linear uptake.

Hence, our derivation yields precise correction factors in the excluded-volume regime *β* = 1, where the ratio of cell-volume and cell-distance is conserved in the limit *ϵ* → 0. Namely, we found that the solution depends on scaling factors *a*(*ρ*) for the diffusion constant and *f*(*ρ*) for the cytokine degradation rate, both depending on the cell-density *ρ*. That finding is in line with the intuition of high cell-densities inhibiting diffusion due to excluded-volume effects, and cell-independent cytokine degradation depending on the effective volume of the diffusion domain.

### 3.3 Numerical verification of *L*^2^ convergence

Under the same assumptions as in Section 3.2 and further assuming *β* = 1, we numerically solved both the rescaled cytokine model (7) with varying *ϵ* values and the homogenized model (9) within a cubic domain Ω (Figure 3a). A single secreting cell is positioned at the center, surrounded by responder cells placed on a cubic grid. We investigated two representative settings: a low cell-density scenario with *ρ* = 0.25 and a high cell-density scenario with *ρ* = 0.45. For each density, the diffusion correction coefficient *a*(*ρ*) was computed numerically as described in Section 3.1.

**Figure 3.**
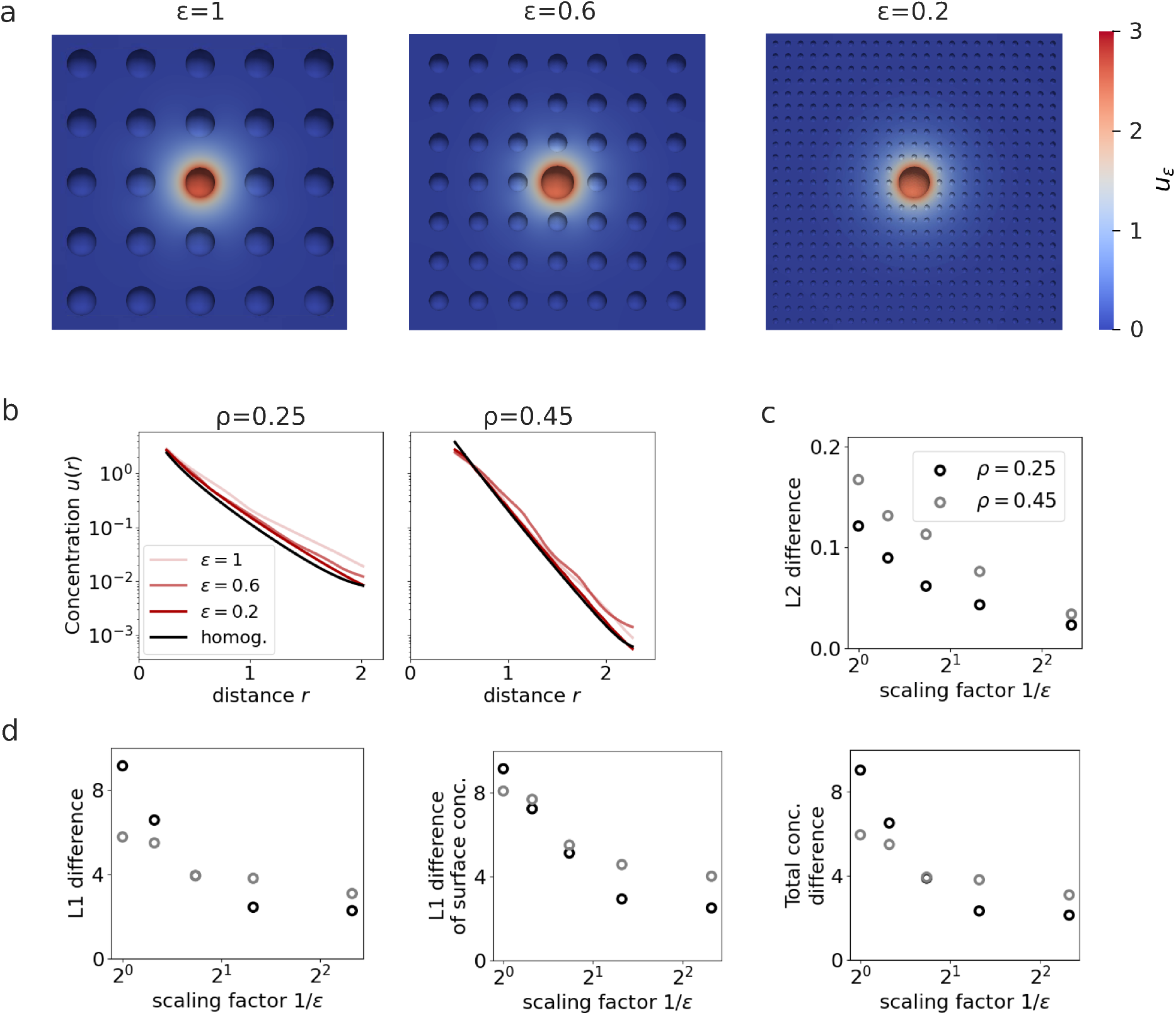
Convergence of the rescaled reaction-diffusion system to the homogenized model. (a) Simulation of the rescaled RD-system (7) with *ϵ* ∈ {1, 0.6, 0.2}. (b) Averaged concentration profile 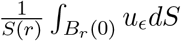in the RD-systems and homogenized model. (c) *L*^2^-difference ||*u*_*ϵ*_ − *u*||*L*_2_ assuming low (*ρ* = 0.25) and high cell density (*ρ* = 0.45). (d) Differences in cytokine concentrations quantified by the *L*^1^-difference ||*u*_*ϵ*_ − *u*||*L*_1_, the summed absolute difference of surface concentrati∫ons at sens∫or cell locations Σ cell *x*_*i*_ |*u*_*ϵ*_(*x*_*i*_) − *u*(*x*_*i*_)|, and the difference in total cytokine amount|∫ _Ω_ *u*_*ϵ*_ *dx* − ∫ _Ω_ *u dx*|.

The spatial cytokine concentration profiles obtained from the rescaled microscopic and homogenized cytokine models exhibit closely matching concentration levels (Figure 3b). This demonstrates that the homogenized formulation provides an accurate approximation of the *ϵ*-rescaled system even at finite *ϵ*. In particular, the *L*^2^-difference between the microscopic solution *u*_*ϵ*_ and the homogenized solution *u* shows the expected vanishing behaviour (Figure 3c-d),

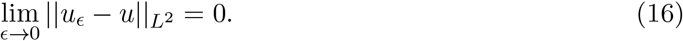

In addition to the *L*^2^-convergence, we evaluated the *L*^1^-norm of the difference and the deviation in total cytokine concentration. Both quantities yield the same vanishing trend as *ϵ* → 0, which was expected as a consequence of the *L*^2^ convergence and the uniform boundedness of the solutions. Furthermore, convergence is observed in both density regimes, thereby validating the homogenized formulation in the excluded-volume limit.

## 4 Cell-population dynamics driven by diffusible ligands

Building on the theoretical findings from the previous sections, we next derived a simulation workflow based on the homogenized description, to analyse cellular populations where cells interact by means of diffusible ligands. We focused specifically on a branching model, where a naive T cell can differentiate into either Th1 or Tfh effector cells, depending on the cytokine environment. Another aim is to assess the model’s accuracy in approximating the full RD system under realistic conditions. Specifically, we evaluated the homogenized model’s performance in predicting cellular activation levels, comparing it directly with the detailed RD model.

First, we sought to to evaluate cellular activation by cytokine signaling in a biological setting (Figure 4a). For that purpose, we first considered a single cytokine secreting cell and placed it on a grid of receptor cells. This configuration mirrors the microscopic geometry studied in the analytical sections. By zooming out, i.e. letting cell size of receptor cells and cell-cell distance converge to 0 at the same speed (case *β* = 1), we arrived at the homogenized cytokine field produced by one secreting cell (Figure 4b). Here, we assumed spherical cells, so that for each secreting cell, the homogenized description of cytokine distribution is given by the radial boundary-value problem (13) derived in Section 3.2.

**Figure 4.**
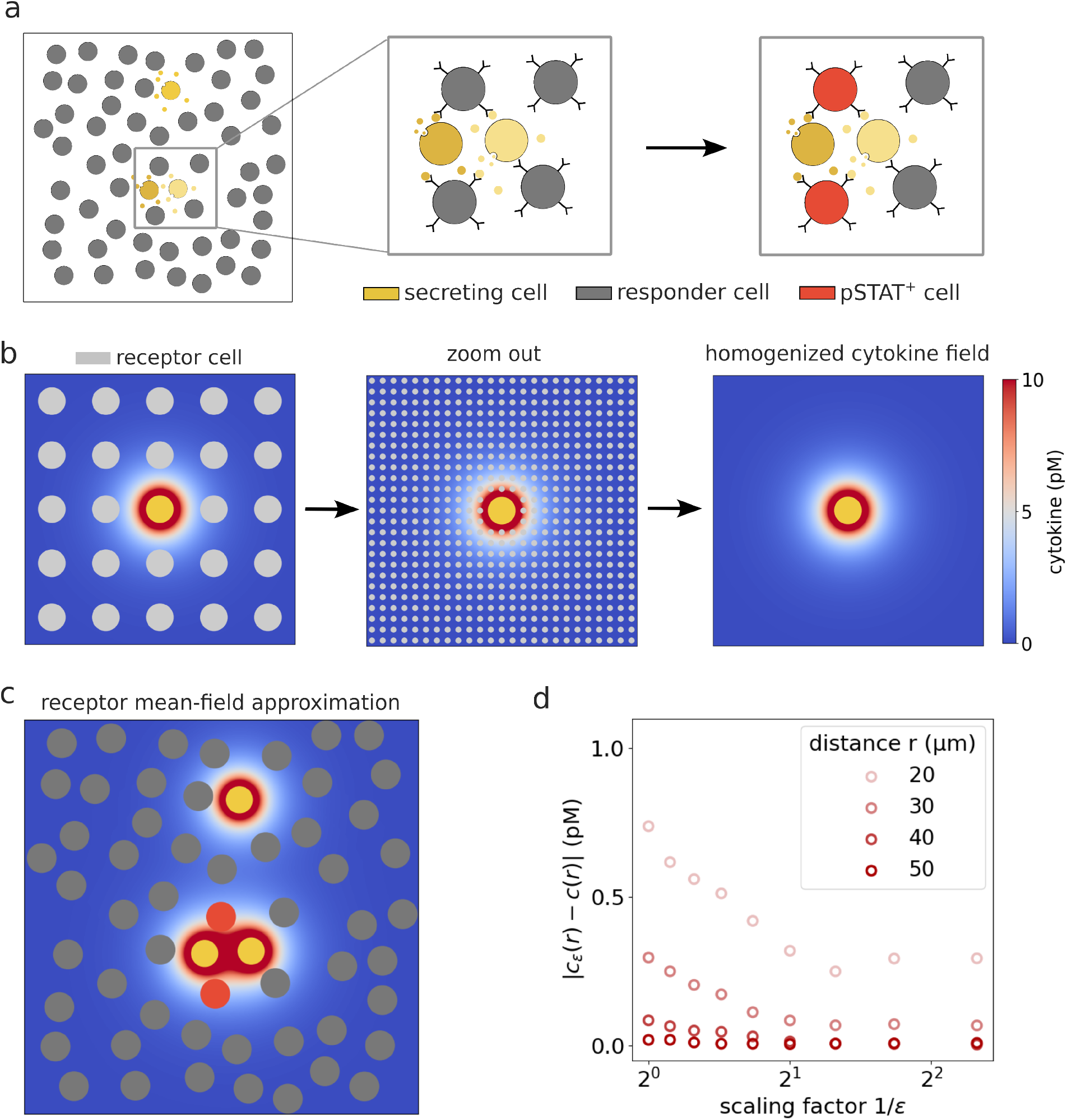
Workflow for evaluating cytokine concentration and responder activation in a biological setting using the homogenized model. (a) Biological setting: cytokine secreting cells (yellow) release cytokines that diffuse through the extracellular space. Responder cells (grey) become activated, that is pSTAT+ (red), when local cytokine concentrations exceed an activation threshold. (b) A single secreting cell is placed on a grid of receptor cells, which are then rescaled and homogenized. (c) Cytokine fields of multiple secreting cells are superposed to evaluate surface concentrations and pSTAT activation at the responder cells. (d) Comparison of the rescaled RD-solution *c*_*ϵ*_ and the homogenized solution *c*, evaluated with physical units.

Then, we returned to the biological setting with multiple secreting cells placed at arbitrary spatial locations. The total cytokine field is obtained by superposing the homogenized cytokine fields associated with each secreting cell, while the effective uptake rate is taken as

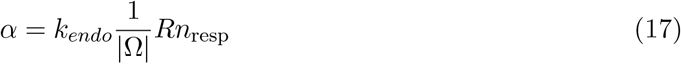

with *n*_resp_ denoting the number of responder cells, and |Ω| the volume of Ω. This expression reflects the homogenized description, in which receptor-mediated uptake is averaged over the population of responder cells.

Next, to evaluate signaling on the single-cell level, cytokine concentrations were evaluated at the surfaces of responder cells (Figure 4c). More precisely, for a responder cell at position *x*, the surface concentration was approximated by averaging the homogenized field over *N*_surf_ = 50 evenly sampled surface points,

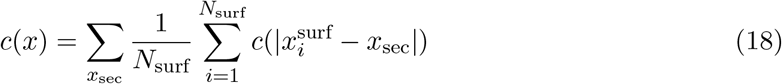

where *x*_sec_ are the positions of the secreting cells and *x*^surf^ are surface points on the responder cell. Following previous work (Brunner et al., 2024), based on the cytokine surface concentration, cellular activation was quantified via the downstream phosphorylation response of the Signal Transducer and Activator of Transcription (pSTAT), modeled by the Hill function 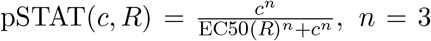,. The cell is considered activated (pSTAT^+^) if its Pstat level has reached a value of pSTAT(*c, R*) ≥ 0.5 (Figure 4c). According to experimental data (Cotari et al., 2013), the effective half-maximal concentration EC50 depends on receptor abundance according to 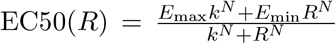, where *E*_max_, *E*_min_, *k* and *N* are fitted parameters (Brunner et al., 2024). To account for receptor heterogeneity, the number of receptors *R* were sampled independently for each responder cell from a log-normal distribution with standard deviation 1.

A direct numerical comparison of the *ϵ*-rescaled RD-system with the homogenized model shows that the concentration profiles closely match, particularly at larger distances to the secreting cell (Figure 4d), While the discrepancy generally decreases with increasing *ϵ*, a slight increase is observed for larger *ϵ* due to limited spatial resolution of the numerical mesh relative to the rescaled cell sizes. Hence, the homogenized cytokine field can be regarded as a close approximation of the finite-*ϵ* setting, further substantiating our workflow.

Using the workflow described above, we systematically evaluated the accuracy of the homogenized model in approximating the full RD model of spatiotemporal cytokine dynamics. For that purpose, both models were simulated using identical physical parameters (Table 1) and the same randomly generated spatial cell configurations (Figure 5a), with prescribed average cell-cell distances. For each configuration, we varied the fraction of cytokine-secreting cells in a biologically relevant range from 2% up to 40%. This allowed us to assess model agreement across different signaling intensities. We compared the models with respect to three key observables: mean surface cytokine concentration at responder cells, spatial variability of concentration levels across the population, and the resulting fraction of activated pSTAT^+^ cells.

**Table 1.**
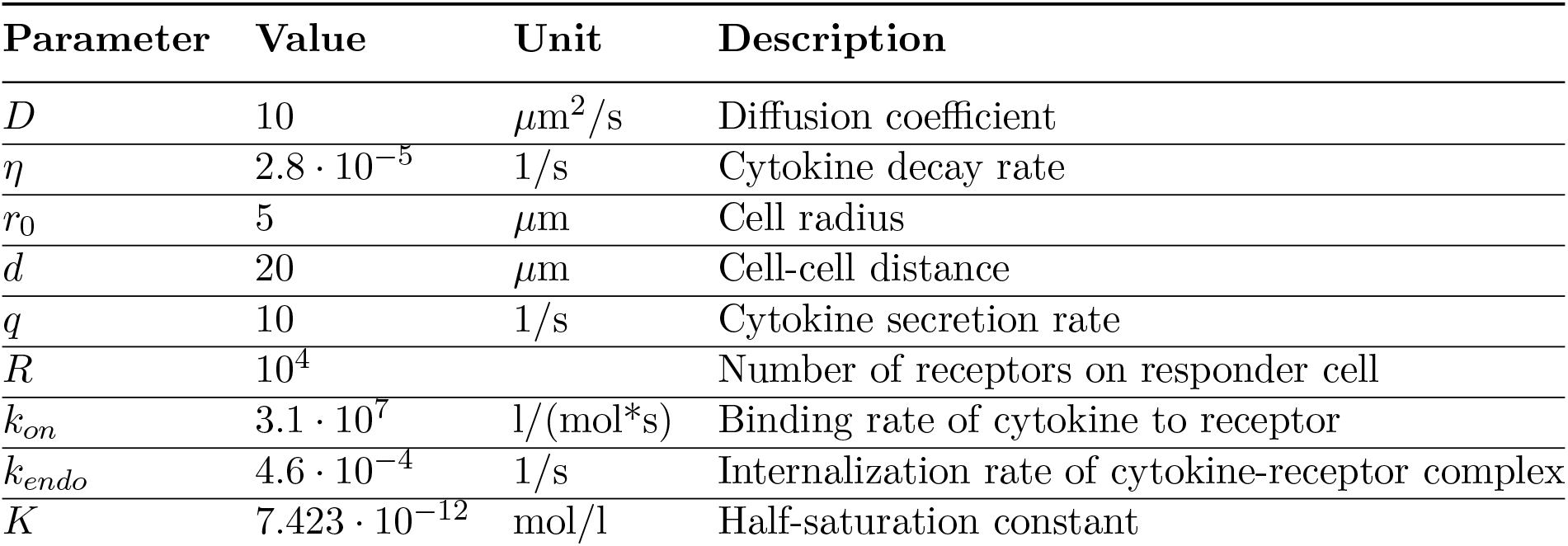
List of physical parameters.

**Figure 5.**
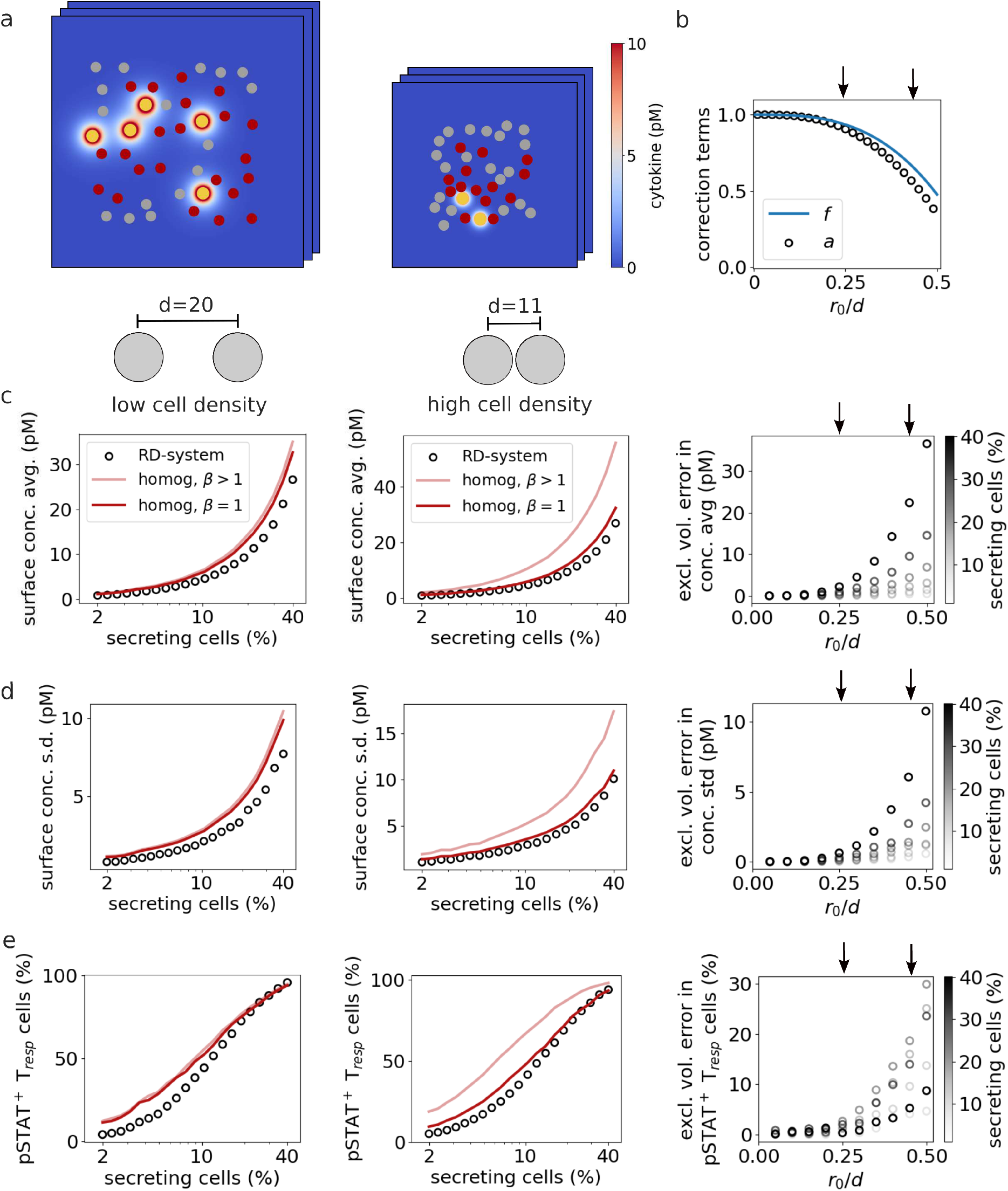
Comparison of the homogenized model and the full reaction-diffusion system under low and high cell density. (a) Simulation visualization. Cells have an average cell-cell distance of 20*µ*m in the low density setting (left) and an average distance of 11*µ*m in the high density setting (right). (b) Correction terms. Shown are the excluded-volume correction terms *f* for degradation and *a* for diffusion depending on the cell density *r*_0_*/d*. (c-e) Model comparison. Surface concentration, spatial standard deviation (s.d.) of cytokine levels across sensor cells and percentage of pSTAT5^+^ cells for varying fractions of cytokine secreting cells, in the RD-system, the homogenized model with *β >* 1 and homogenized model with *β* = 1. Left panel shows the low density setting and middle panel the high density setting. Right panel shows the excluded volume error, i.e. the difference between the homogenized models with *β >* 1 and *β* = 1.

In a low cell-density scenario with average cell-cell distance *d* = 20*µm* and cell density 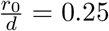, the excluded-volume correction factors for diffusion and degradation are close to 1 (Figure 5b). Therefore, the discrepancy between the homogenized models corresponding to the scaling regimes *β >* 1 and *β* = 1 are small, and both closely match the precise RD model across all observables (Figure 5c-e, left panel). In contrast, at higher cell densities with average cell-cell distance *d* = 11*µm* and cell density 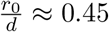, excluded-volume effects become significant. Here, the homogenized model derived under the scaling *β* = 1, which incorporates effective diffusion and degradation corrections, aligns well with the RD-system. However, the model corresponding to *β >* 1, which neglects these corrections, shows substantial deviations in predicted concentrations and activation levels (Figure 5c-e, middle panel). To quantify this discrepancy, we evaluated the excluded volume error, defined as the difference between the homogenized solutions obtained with *β* = 1 and *β >* 1. We observed that the error increases monotonically with the volume fraction *r*_0_*/d*, confirming that crowding-induced modifications become increasingly important as the cell volume fraction grows (Figure 5c-e, right panels).

Next, we applied our workflow to study a cytokine-mediated branching model (Figure 6a), illustrating how the homogenization approach can be used to efficiently evaluate cytokine fields in more complex cellular scenarios. In this model, a naive Th cell differentiates into either a Tfh or Th1 effector cell with differentiation probabilities *p*_Tfh_ and *p*_Th1_ respectively. The probabilities are determined by norming weighted Hill-type functions of the local concentrations *c* of IL-21 and 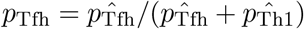 and 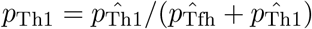, where

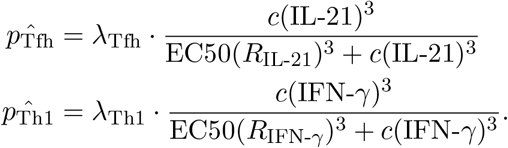

**Figure 6.**
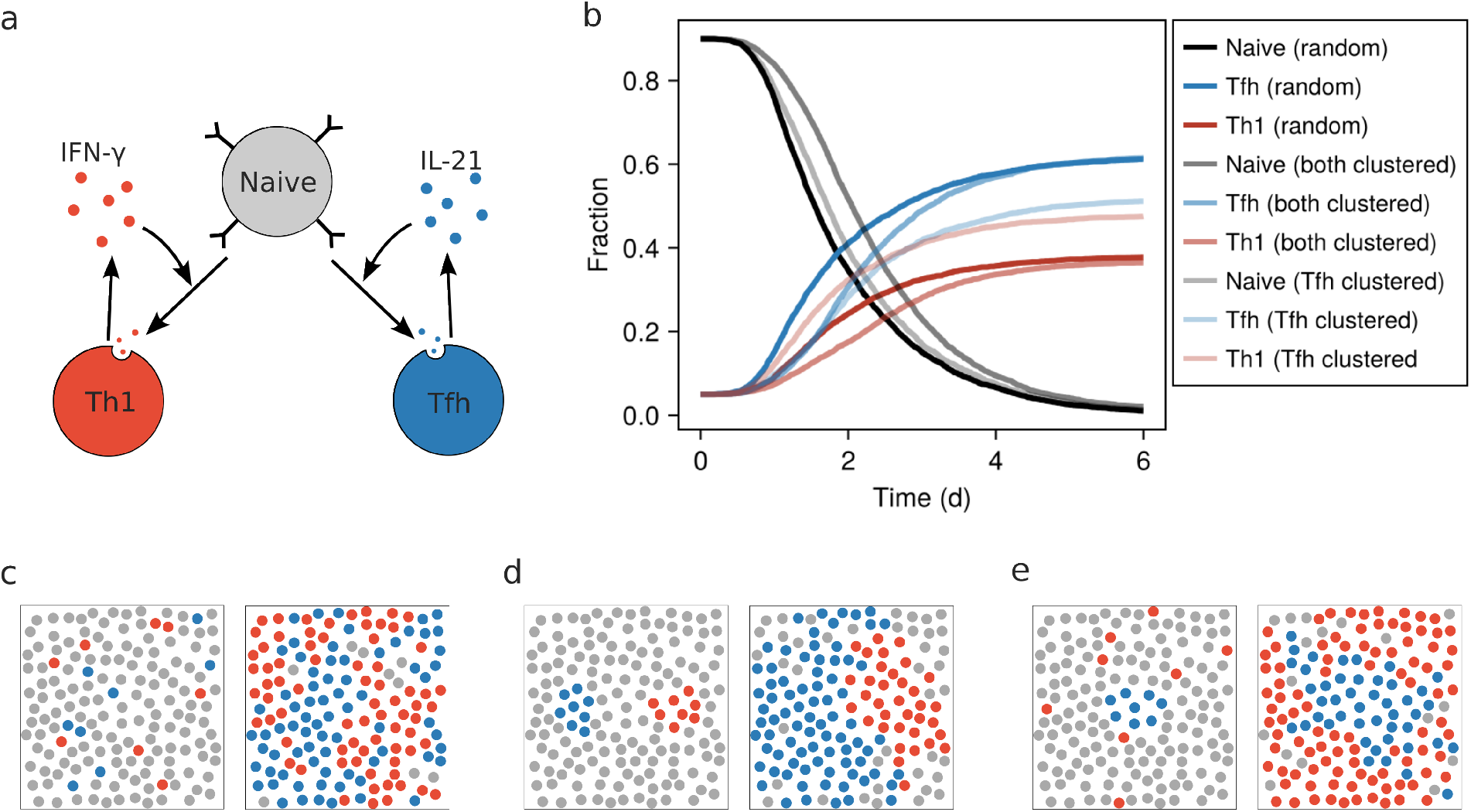
Influence of local cytokine signaling and initial cell positions on branching differentiation. (a) Model scheme. A naive cell with receptor count *R*_IL-21_ = *R*_IFN-*γ*_ = 1500 differentiates into either Th1 or Tfh effector cell. The decision depends on the local concentrations of IL-21 and IFN-*γ*, which are secreted by the effector cells. Differentiation weights are chosen to be *λ*_Tfh_ = 1.2 and *λ*_Th1_ = 0.8, and the differentiation process is delayed by the gamma distribution with shape-parameter 5 and rate-parameter 0.003 min^−1^. (b) Kinetics of the total fraction of differentiated cells for random versus initially clustered effector cell configurations. (c-e) Spatial distribution of differentiated cells at time point 0d (left) and 6d (right) for random initial positioning (c), clustered initial positioning of both effector cells (d) and clustered initial positioning of Tfh cells only (e).

Cytokine concentrations at the naive cells are computed using the homogenized cytokine field, thus extending our previous work where a similar system was studied using ODE formulations neglecting spatial effects (Burt and Thurley, 2023). Differentiation is implemented as a stochastic process with gamma-distributed waiting times (Thurley et al., 2018), introducing a delay between cytokine exposure and final differentiation. Once differentiated, effector cells secrete their corresponding cytokine, thereby establishing positive feedback loops: Tfh cells reinforce Tfh differentiation via IL-21, and Th1 cells analogously promote Th1 differentiation via IFN-*γ*. Cytokine fields are assumed to be in quasi-steady state relative to the differentiation dynamics, consistent with the timescale separation underlying the homogenized model.

Starting from an initial condition in which 5% of cells are assigned to each effector cell type, the fractions of Tfh and Th1 cells increase steadily over time until all cells are differentiated. While the overall differentiation kinetics are governed by cytokine production rates and receptor sensitivity, the spatial organization of initial effector cells substantially influences the temporal evolution. When both effector cells are initially clustered, the expansion of both populations is slowed compared to spatially dispersed initial conditions, indicating that clustering reduces the effective range of paracrine signaling (Figure 6b). We additionally considered an asymmetric configuration in which only Tfh cells were initially clustered, whereas Th1 cells were randomly distributed. In this case, not only the differentiation kinetics but also the steady-state fractions were altered. The final proportion of Tfh cells was reduced compared to symmetric or fully random initial conditions. This demonstrates that strong spatial clustering can reduce the overall differentiation into the clustered cell type, as cytokine-mediated recruitment of distant naive cells is restricted. Hence, spatial confinement does not only delay expansion but can shift the long-term population balance.

Next, we examined the spatial organization of the cell populations over time. Even when effector cells are initially distributed randomly, local cytokine reinforcement promotes the differentiation of nearby naive cells into the same effector type, resulting in the spontaneous formation of spatial local clusters (Figure 6c). When effector cells are initially arranged in clusters, this mechanism is further strengthened, leading to pronounced spatial segregation of Tfh and Th1 cells that reflects and reinforces the initial configuration (Figure 6d). In the asymmetric scenario where only Tfh cells are initially clustered, the Tfh cluster expands over time, whereas Th1 cells emerge in the surrounding regions (Figure 6e).

Overall, homogenizing over the responder cells yielded a modeling workflow that efficiently captures cytokine concentration profiles and cellular activation levels. Coupling to a branching model of T cell differentiation revealed how spatial organization influences both transient dynamics and steady-state population composition.

## 5 Discussion

The aim of this study was to develop a rigorous and tractable approach for describing cytokine-mediated communication in large and geometrically complex cell populations. While the microscopic formulation of the RD system provides a detailed mechanistic model, its computational cost grows rapidly with the number of interacting cells. On the other hand, existing explicit approximations such as Yukawa-type solutions lack a precise connection to the underlying cellular geometry. Applying homogenization techniques results in a macroscopic model that replaces discrete cells with a continuous density while retaining essential aspects of cellular uptake and excluded volume. Of note, the homogenization process does not impose any constraints with respect to the size and shape of cytokine-secreting cells, which therefore can be chosen in line with the application. In case of secreting cells with radial symmetry, the macroscopic formulation further simplifies to an ODE system or even explicit formulas describing cytokine gradients, thus allowing for numerically highly efficient integration into scalable models of cell-population dynamics.

Our results indicate that finite cell volumes can have a substantial effect on cytokine transport, since high cell densities inhibit effective diffusion and degradation, leading to correction factors in the homogenized limit. This explains why, in the excluded-volume regime *β* = 1, the macroscopic equation does not reduce to the classical screened Poisson equation, and therefore does not admit a pure Yukawa potential solution. However, as the cell-volume fraction tends to zero, the excluded-volume factors converge to 1, continuously linking the two regimes *β* = 1 and *β >* 1. Hence, these effects are particularly relevant when modelling densely packed tissues such as lymph nodes or the tumor microenvironment.

As an illustrative example, we used our approach to analyze a Th branching model in a spatially resolved setting. Although Th subset differentiation has been extensively investigated in reductionist in vitro systems using recombinant cytokines, the mechanisms governing this process in vivo are considerably more complex and only beginning to be understood. Experimentally, the analysis of Th subset differentiation in vivo is challenging due to the very low frequencies of antigen-specific T cells during the early stages of an immune response and the difficulty of accurately determining the cytokine concentrations experienced by individual T cells within lymphoid tissues. In vivo, local cytokine availability is shaped not only by the identity and spatial distribution of cytokine-producing cells but also by the presence and localization of cytokine-consuming cells. In addition, cytokine diffusion within tissues is constrained by tissue architecture and microenvironmental structure. Together, this results in spatially restricted cytokine niches within lymphoid tissues. Large-scale in silico simulations can substantially help to understand the effect of different cytokines on Th subset differentiation in vivo. The homogenization approach eliminates the need to resolve individual cell positions and provides an explicit link between microscopic parameters (cell size, cell-cell distance) and macroscopic observables (effective diffusivity and degradation). This makes the approach especially suitable for multi-cellular scenarios involving long-range communication or multiple cytokine species.

Spatial cytokine concentrations can vary over several orders of magnitude, even when only a single diffusible species is involved and diffusion operates on a much faster timescale than cellular secretion and uptake. This naturally prompts the question of how such structured fields relate to classical mechanisms of pattern formation in RD systems. In traditional Turing-type settings, spatial patterns arise from instabilities generated by interactions between at least two chemical species with different diffusivity(Klika et al., 2011; Marcon and Sharpe, 2012). The cytokine fields considered here behave differently. They do not produce de novo periodic patterns from homogeneous initial conditions. Nevertheless, model simulations revealed that a single cytokine species can generate stable spatial profiles even under quasi-steady-state conditions (Oyler-Yaniv et al., 2017; Thurley et al., 2015), and the strict spatial localization of cytokine sources and sinks gives rise to cytokine gradients spanning several orders of magnitude (Brunner et al., 2024). The resulting heterogeneity therefore reflects positional information shaped by geometric organization and cell-tissue architecture rather than reaction-driven instabilities (Raspopovic et al., 2014; Wolpert, 1969). Similar behavior has been observed in various biological systems, where spatial organization is influenced by localized cell-cell interactions and geometric factors, such as in models of cell migration and cancer metastasis (Marciniak-Czochra et al., 2017; Mensah et al., 2025; Painter, 2019).

While the homogenized model captures key features of cytokine dynamics, it relies on several simplifying assumptions. In particular, the current model assumes homogeneous secretion and uptake across each cell surface and a uniform cell shape of responder cells. In lymphoid tissues, receptor expression and cell morphology may vary, potentially altering local cytokine gradients in nontrivial ways. Extending the approach to incorporate spatial heterogeneity in cell properties and tissue architecture therefore represents an important and biological relevant direction for future work. For example, existing studies have coupled signaling processes to cell mechanics, providing a framework to study the impact of cell shape on signaling dynamics (Verhees et al., 2025).

Overall, by linking microscopic biophysical rules to macroscopic cytokine fields, the homogenized model reveals how individual cellular properties collectively determine cytokine gradients. Its integration with models of immune cell motility and differentiation promises deeper insights into how spatial organization, tissue structure and cell density shape immune-cell decision-making across scales.

## Appendix A Proof of Proposition 1

We consider the rescaled PDE (7) with uptake function Ψ(*u*) = *u* in the case of linear uptake, and 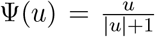in the case of saturated uptake. There exists a unique weak solution *u*_*ϵ*_ in *W*_*ϵ*_ := {*w* ∈ *W* ^1,2^(Ω_*ϵ*_) : *w* = 0 on (∂Ω)_0_} and we determine its limit in *W* := {*w* ∈ *W* ^1,2^(Ω) : *w* = 0 on (∂Ω)_0_} in the following.

### Step 1: Extension of *u*_*ϵ*_ to *W*

By testing the variational formulation of (7),

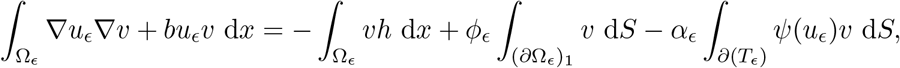

with *u*_*ϵ*_, we obtain a uniform bound on ∥*u*_*ϵ*_ ∥ _*W*_*ϵ*. By classical extension results (Cioranescu and Paulin, 1979; Conca and Donato, 1988), there exists a constant *c >* 0 independent of *ϵ* and a family of linear and bounded operators *P*_*ϵ*_ : *W*_*ϵ*_ → *W* such that for all *w* ∈ *W*_*ϵ*_

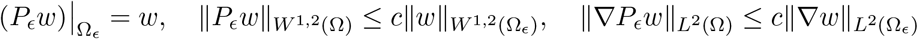

Thus, {*P*_*ϵ*_*u*_*ϵ*_} is uniformly bounded in *W*, and therefore *P*_*ϵ*_*u*_*ϵ*_ ⇀ *u* in *W* for a subsequence. Since the limit *u* is unique and therefore independent of the choice of the extension, we write *u*_*ϵ*_ for its extension *P*_*ϵ*_*u*_*ϵ*_ in the following. It remains to identify *u*.

### Step 2: Limit of linear terms

When *β >* 1, strong convergence of the characteristic function *χ*_Ω_ε → 1 in *L*^2^(Ω) implies

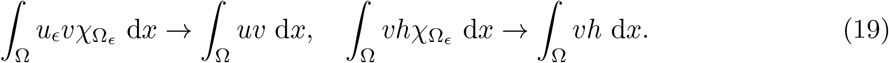

In the case *β* = 1, we only have 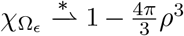 in *L* ∞ (Ω). Since by Sobolev embedding *u*_*ϵ*_ ⇀ *u* in *W* (Ω) implies *u*_*ϵ*_ → *u* in *L*^2^(Ω) strongly, we again have

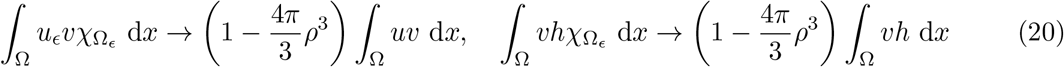

### Step 3: Limit of the (non-linear) uptake term

Let *β* ∈ [1, 3), *v* ∈ *C*^1^(Ω), and consider non-linear uptake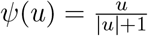. It holds that

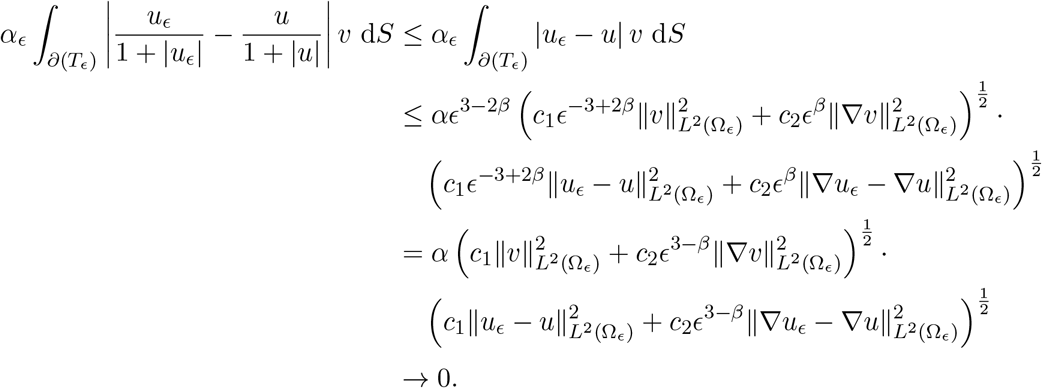

Of note, the last expression converges to 0 since we assumed *β <* 3. Thus, it suffices to determine the limit of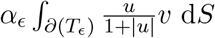.

By approximation, it is sufficient to consider *α*_*ϵ*_ *∂*(*T*_*ϵ*_) *φ* d*S* for *φ* ∈ *C*(Ω). Take for *ϵ* an enumeration *T*_*ϵ,i*_ of the holes comprising the set ∂(*T*_*ϵ*_) and denote by *x*^*ϵ*^ the minimum of *φ* on *T*_*ϵ,i*_. Then

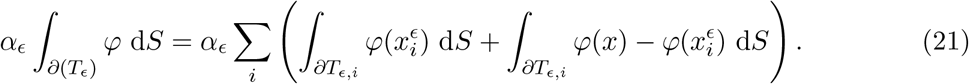

Since *r*(*ϵ*) = *ρϵ*^*β*^ we have

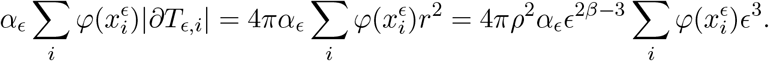

It holds that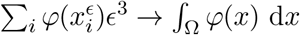, so

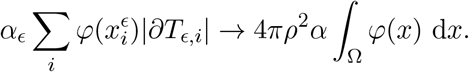

The second part of (21) can be estimated by

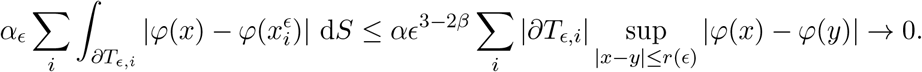

All in all, we have

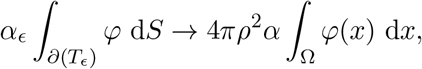

By approximation, we conclude

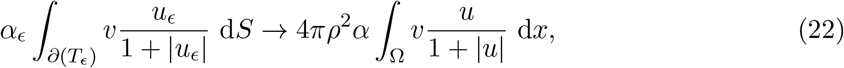

for *v* ∈ *W*.

By the same proof, it holds

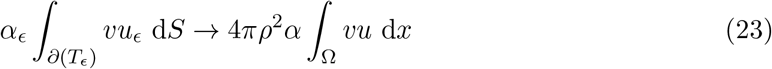

for linear uptake and *β* ∈ [1, 3). The proof for linear Neumann boundary conditions are also considered in the classical paper (Cioranescu and Donato, 1988).

### Step 4: Limit of the diffusion term

When *β >* 1, *χ*_Ω_ε → 1 in *L*^2^(Ω) implies

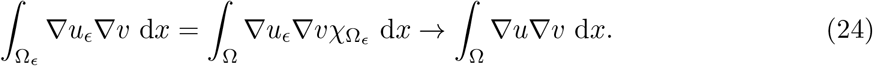

For *β* = 1, it only holds 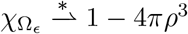in *L*^∞^(Ω), which does not imply (24). It is a classical result that instead

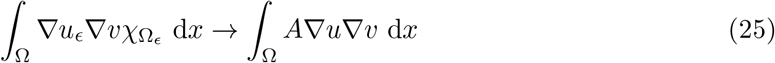

for *A* defined in Proposition 1, for the proof see e.g. (Cioranescu and Donato, 1988; Cioranescu and Murat, 2018).

## APPENDIX B Software and simulations

Spatial simulations of the cell problem (10) (cf. Figure 2) were carried out using FEniCSx with first-order Lagrange finite elements. The extracellular domain was discretized with tetrahedral meshes generated in Gmsh. To accurately resolve boundary layers near the cell surface, mesh refinement was controlled by a distance-based field centered at the cell boundary. Periodic boundary conditions were enforced using multi-point constraints, implemented via the DOLFINx-MPC package. Simulations of the rescaled RD system (cf. Figure 3) were performed using a modified version of the FEniCS-based software developed in Brunner et al. (2024). Cells were positioned on a rescaled three dimensional grid using custom locator functions, with a single secreting cell at the domain center. For convergence validation, the homogenized spherical boundary-value problem (13) was solved with SciPy’s boundary-value problem solver. The resulting radial solution was then evaluated at corresponding spatial locations and projected onto the finite-element mesh of the rescaled RD-system to enable direct pointwise comparison. The spatial simulations used for model comparison (cf. Figure 5) were carried out using the same FEniCS-based software. Cell positions were generated using a Bridson Poisson-disk sampling algorithm to ensure a prescribed minimal cell-cell distance while maintaining spatial randomness. For each configuration, secreting cells were randomly selected from the fixed cell positions according to the specified fraction. Each parameter setting was simulated in 20 independent realizations, and reported quantities were averaged over these runs. The branching model was implemented in Python using modular components for parameter input, initialization, cytokine field evaluation, differentiation dynamics, and data output. Simulations were initialized from a text-based experiment file that specifies cell-type parameters and cell positions which are given as precomputed Bridson Poisson-disk point sets. At each time step, cytokine concentrations at responder cell surfaces were computed by superposition of the homogenized ODE solution (5) at producer cell positions.

